# What do ossification sequences tell us about the origin of extant amphibians?

**DOI:** 10.1101/352609

**Authors:** Michel Laurin, Océane Lapauze, David Marjanović

**Author notes:** **Cite as:** Laurin M, Lapauze O, and Marjanović D (2019). What do ossification sequences tell us about the origin of extant amphibians? *bioRxiv* 352609, ver. 4 peer-reviewed by PCI Paleo. DOI: 10.1101/352609.

## Abstract

The origin of extant amphibians has been studied using several sources of data and methods, including phylogenetic analyses of morphological data, molecular dating, stratigraphic data, and integration of ossification sequence data, but a consensus about their affinities with other Paleozoic tetrapods has failed to emerge. We have compiled five datasets to assess the relative support for six competing hypotheses about the origin of extant amphibians: a monophyletic origin among temnospondyls, a monophyletic origin among lepospondyls, a diphyletic origin among both temnospondyls and lepospondyls, a diphyletic origin among temnospondyls alone, and two variants of a triphyletic origin, in which anurans and urodeles come from different temnospondyl taxa while caecilians come from lepospondyls and are either closer to anurans and urodeles or to amniotes. Our datasets comprise ossification sequences of up to 107 terminal taxa and up to eight cranial bones, and up to 65 terminal taxa and up to seven appendicular bones, respectively. Among extinct taxa, only two or three temnospondyl can be analyzed simultaneously for cranial data, but this is not an insuperable problem because each of the six tested hypotheses implies a different position of temnospondyls and caecilians relative to other sampled taxa. For appendicular data, more extinct taxa can be analyzed, including some lepospondyls and the finned tetrapodomorph *Eusthenopteron*, in addition to temnospondyls. The data are analyzed through maximum likelihood, and the AICc (corrected Akaike Information Criterion) weights of the six hypotheses allow us to assess their relative support. By an unexpectedly large margin, our analyses of the cranial data support a monophyletic origin among lepospondyls; a monophyletic origin among temnospondyls, the current near-consensus, is a distant second. All other hypotheses are exceedingly unlikely according to our data. Surprisingly, analysis of the appendicular data supports triphyly of extant amphibians within a clade that unites lepospondyls and temnospondyls, contrary to all phylogenies based on molecular data and recent trees based on paleontological data, but this conclusion is not very robust.

## Introduction

Paleontologists have been studying the origin of the extant amphibian clades for more than a century. Early studies generally proposed an origin of at least some extant amphibians from temnospondyls. **Cope (1888)** initially suggested that batrachians (anurans and urodeles) derived from temnospondyls (a large clade of limbed vertebrates known from the Early Carboniferous to the Early Cretaceous) because he believed that the batrachian vertebral centrum was an intercentrum, the dominant central element of temnospondyls. Later, **Watson (1940)** argued that anurans were derived from temnospondyls because of similarities (mostly in the palate) between the temnospondyl “*Miobatrachus*” (now considered a junior synonym of *Amphibamus*) and anurans. Monophyly of extant amphibians (Lissamphibia) was proposed by **Parsons and Williams (1962, 1963)**, an idea that was accepted more quickly by herpetologists than by paleontologists. Lissamphibian monophyly was supported by (among a few other character states) the widespread occurrence of pedicellate, bicuspid teeth. The subsequent discovery of such teeth in the amphibamid temnospondyl *Doleserpeton* (**Bolt, 1969**) reinforced the widespread acceptance of an origin of Lissamphibia from within temnospondyls (e.g., **Schoch and Milner, 2004**). Recently, this hypothesis, referred to as the temnospondyl hypothesis or TH for short (**Fig. 1c**), has been supported by several phylogenetic analyses based on phenotypic data matrices (e.g., **Ruta and Coates, 2007**; **Sigurdsen and Green, 2011**; **Maddin et al., 2012**; **Pardo et al., 2017a, fig. S6**; **Pardo et al., 2017b**; **Mann et al., 2019**).

**Figure 1.**
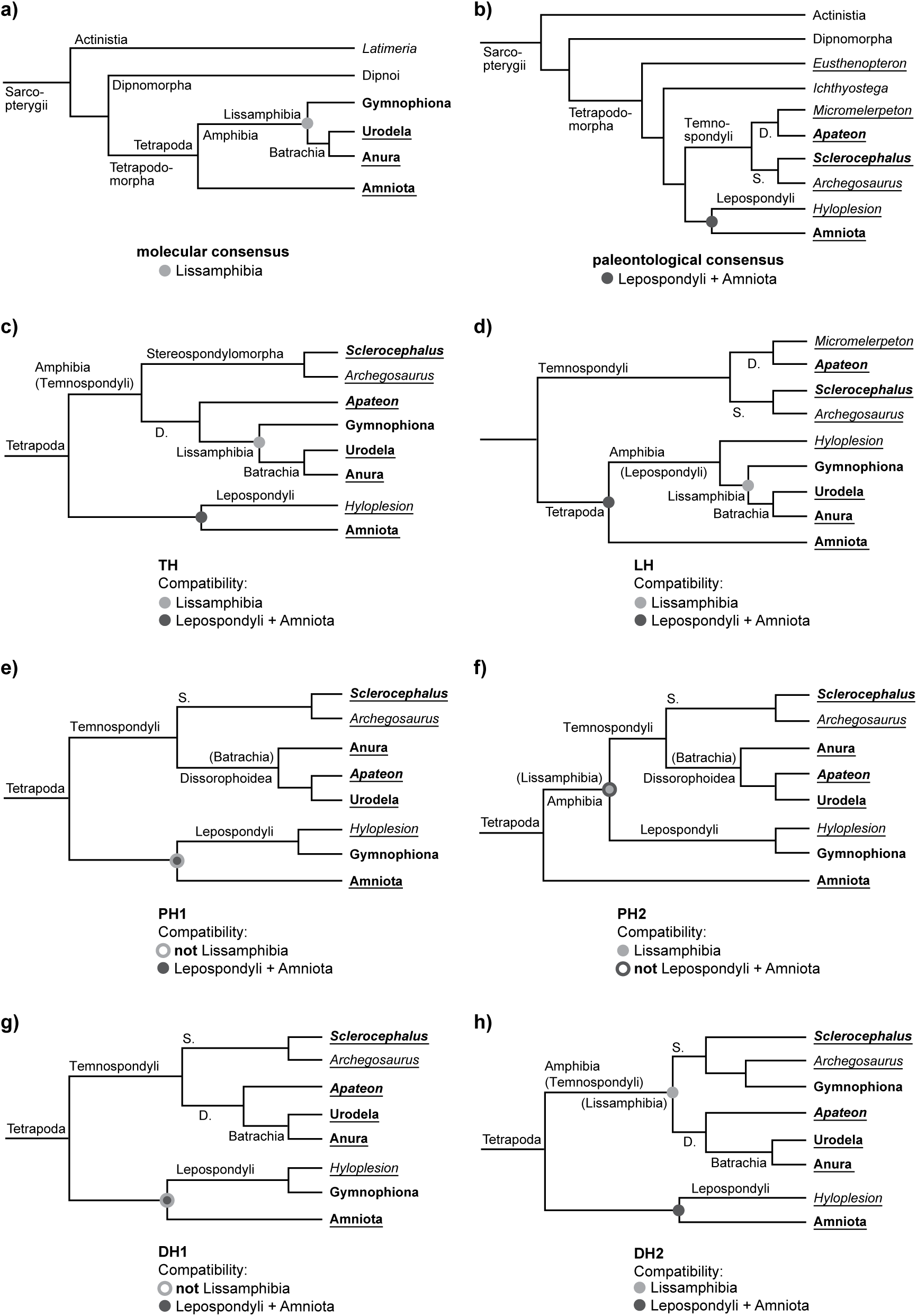
Hypotheses on the relationships of the extant amphibian clades since the late 20th century. The names of terminal taxa sampled here for cranial characters are in boldface, those sampled for appendicular characters are underlined; the names of larger clades are placed toward the right end of a branch if they have minimum-clade (node-based) definitions, to the left if they have maximum-clade (branch-based) definitions. Names in parentheses would, given that phylogenetic hypothesis, not be used, but replaced by synonyms. Among terminal taxa, “*Melanerpeton*” *humbergense*, sampled for appendicular characters, is not shown, but is always the sister-group of *Apateon*; *Microbrachis*, likewise sampled for appendicular characters, is not shown either, but is always the sister-group of *Hyloplesion*; *Eusthenopteron* is not shown in c)–h), where it forms the outgroup (b)). See text for *Micromelerpeton* and for references. The first two trees (a, b) show the current consensus; the other trees (c–h) show the various tested paleontological hypotheses. *Abbreviations:* D., Dissorophoidea; S., Stereospondylomorpha. **a)** Consensus of the latest phylogenetic analyses of molecular data; all named clades are therefore extant. Note the monophyly of the extant amphibians (Lissamphibia, marked with a light gray dot) with respect to Amniota. **b)** Consensus of all recent analyses of Paleozoic limbed vertebrates, omitting the extant amphibian clades. Note the monophyly of “lepospondyls” + amniotes (marked with a dark gray dot). **c)** TH: “temnospondyl hypothesis”. Lissamphibia nested among dissorophoid temnospondyls. Compatible with both a) and b) (gray dots). **d)** LH: “lepospondyl hypothesis”. Lissamphibia nested among “lepospondyls”; consequently, temnospondyls are not crown-group tetrapods. Compatible with both a) and b) (gray dots). **e)** PH1: “polyphyly hypothesis”, first variant. Urodela as dissorophoid temnospondyls close to *Apateon*, Anura as a separate clade of dissorophoid temnospondyls, Gymnophiona as “lepospondyls”. Compatible with b) (dark gray dot) but not with a) (light gray circle). **f)** PH2: “polyphyly hypothesis”, second variant. Like PH1, but with restored monophyly of extant amphibians with respect to amniotes (light gray dot; see a)) at the expense of compatibility with the paleontological consensus concerning the position of temnospondyls, lepospondyls, and amniotes (dark gray circle; see b)). **g)** DH1: “diphyly hypothesis”, first variant. Batrachia as dissorophoid temnospondyls, Gymnophiona as “lepospondyls”. Compatible with b) (dark gray dot) but not with a) (light gray circle). **h)** DH2: “diphyly hypothesis”, second variant. Batrachia as dissorophoid temnospondyls, Gymnophiona as stereospondylomorph temnospondyls. Compatible with both a) and b).

Other hypotheses about the origin of extant amphibians have been available in the literature for nearly as long a time (see **Schoch and Milner, 2004** for a historical review). These were initially formulated especially for the urodeles and caecilians, which are less similar to temnospondyls and lack a tympanic middle ear (which is present in most anurans and often inferred for at least some temnospondyls but absent in lepospondyls). Thus, **Steen (1938)** highlighted similarities in the palate (broad cultriform process of the parasphenoid) and cheek (loss of several bones) between lysorophian lepospondyls and urodeles. **Carroll and Currie (1975)** and **Carroll and Holmes (1980)** argued that the extant amphibians had three distinct origins among early stegocephalians; while they accepted an origin of anurans among temnospondyls, they suggested that urodeles and caecilians originated from two distinct groups of lepospondyls (*Rhynchonkos* for caecilians, Hapsidopareiidae for urodeles). Later, based mostly on developmental similarities between the temnospondyl *Apateon* and urodeles, **Carroll (2001, 2007)** and **Fröbisch et al. (2007)** proposed another hypothesis involving a triphyletic origin of lissamphibians, with an origin of anurans and urodeles from two distinct temnospondyl groups, while the caecilians would remain in the lepospondyl clade. This is what we call the polyphyly hypothesis (PH). We have tested two versions. One (here called PH1; **Fig. 1e**) was cautiously suggested by **Fröbisch et al. (2007)**; it agrees with the paleontological consensus in placing all or most lepospondyls closer to Amniota than to Temnospondyli (**Fig. 1b**; **Sigurdsen and Green, 2011**; **Pardo et al., 2017a, fig. S6**; **Pardo et al., 2017b**; **Marjanović and Laurin, 2019**; **Clack et al., 2019**; **Mann et al., 2019**). The other (PH2; **Fig. 1f**) is modified to make Lissamphibia monophyletic with respect to Amniota, a fact we consider demonstrated beyond reasonable doubt by multiple phylogenetic analyses of molecular data (**Fig. 1a**; **Irisarri et al., 2017**; **Feng et al., 2017**; and references cited therein); this comes at the expense of contradicting the paleontological consensus, which was not yet established when **Milner (1993, 16–18, fig. 5B)** argued for something like the PH2 as one of two more or less equal possibilities. **Anderson (2007)** and **Anderson et al. (2008)** found lissamphibian diphyly, specifically a monophyletic, exclusive Batrachia among the temnospondyls while keeping the caecilians among the lepospondyls (DH1; **Fig. 1g**). **Pardo et al. (2017a, fig. 2, S7)** presented a similar hypothesis, with batrachians and caecilians having separate origins within the temnospondyls (DH2; **Fig. 1h**); we should point out, however, that their dataset contained only temnospondyls and lissamphibians, and while they found the DH2 using Bayesian inference, it was only one of four equally parsimonious results (see **Marjanović and Laurin, 2019** for this fact and a discussion of Bayesian analysis of paleontological datasets). Further, a monophyletic origin of all extant amphibians among lepospondyls has also been proposed (**Laurin, 1998**; **Pawley, 2006, appendix 16**; **Marjanović and Laurin, 2009, 2013, 2019**). This will be referred to below as the lepospondyl hypothesis (LH; **Fig. 1d**).

**Figure 2.**
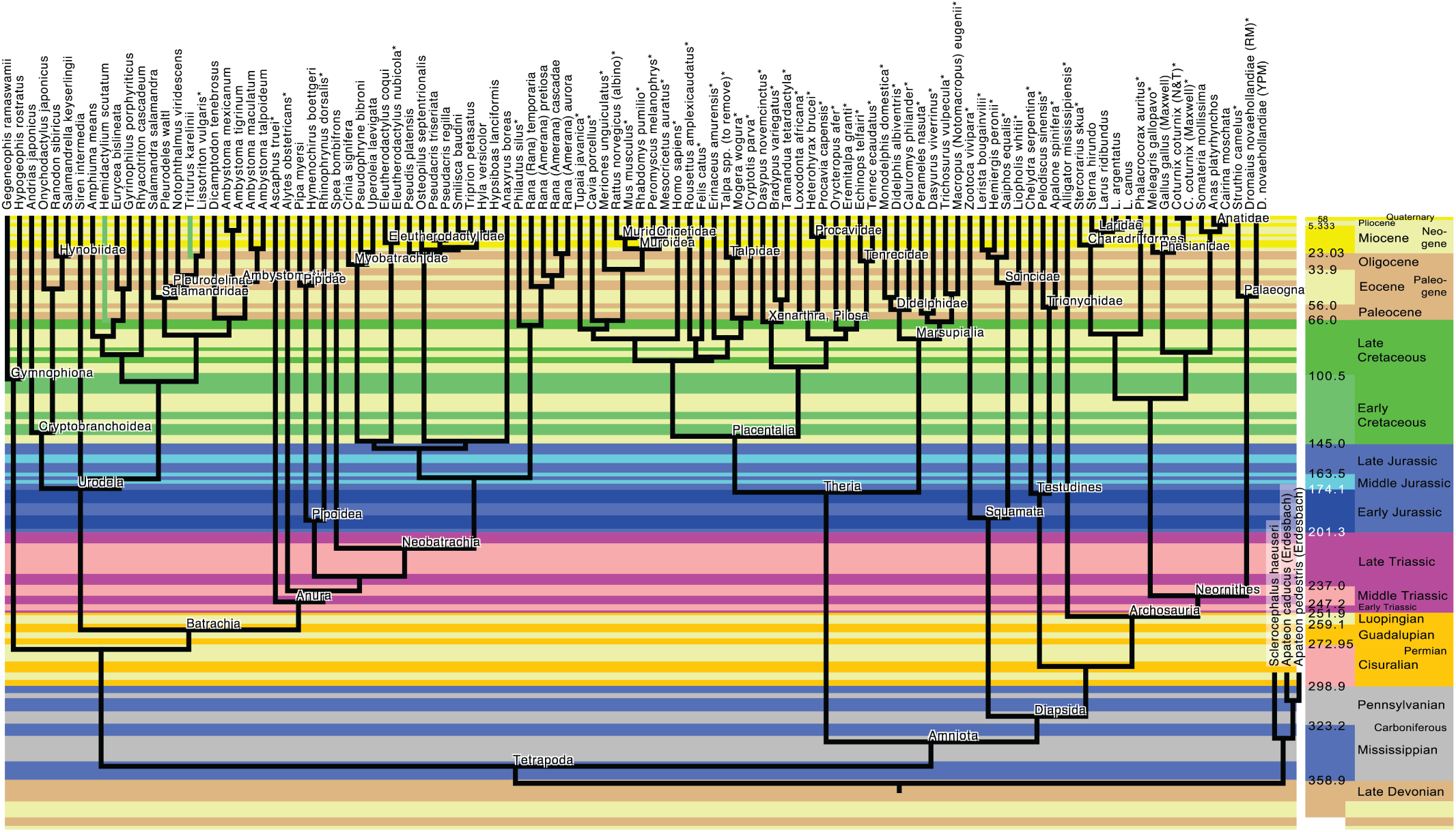
Reference phylogeny used for some of the analyses, illustrating the LH (lepospondyl hypothesis) of lissamphibian origins. The tree was time-calibrated, but analyses showed that branch lengths are irrelevant, given that the best model is speciational (**Tables 2–4**). Main sources for topology and divergence times: **Reeder (2003)**; **Brandley et al. (2005)**; **Pons et al. (2005)**; **Lecompte et al. (2008)**; **Bossuyt and Roelants (2009)**; **Germain and Laurin (2009)**; **Hugall et al. (2007)**; **Gonzalez et al. (2009)**; **Meredith et al. (2011)**; **Sterli et al. (2013)**; **Wang et al. (2013)**; **Marjanović and Laurin (2014, 2019)**); **Pyron (2014)**; **Rabosky et al. (2014)**; **Schoch (2014b)**; **Prum et al. (2015)**; **Zhuang et al. (2015)**; **Tarver et al. (2016)**; **Feng et al. (2017)**; **Irisarri et al. (2017)**; **Lu et al. (2017)**; **Pardo et al. (2017a)**; **Jetz and Pyron (2018)**. The colored bands represent geological stages from the international geological timescale (**Ogg et al., 2016**).

Phylogenetic analyses of molecular data cannot distinguish the TH, the PH2, the DH2 or the LH from each other by topology (**Fig. 1**) because all of these imply lissamphibian monophyly with respect to amniotes, and molecular data are not available from any other tetrapodomorphs. Several other types of data and methods have, however, been used to try to discriminate between the various hypotheses on the origin of extant amphibians. In addition to classical phylogenetic analyses of morphological data matrices, these include the use of molecular dating (**Zhang et al., 2005**; **Marjanović and Laurin, 2007**; **Pardo et al., 2017a**) and stratigraphic data (**Marjanović and Laurin, 2008**) to compare the inferred divergence dates between the three main extant amphibian clades on the basis of molecular data with predictions based on the fossil record under the TH and the LH on one side and the PH and the DH on the other. However, developmental data, in the form of ossification sequences, have been the second-most frequently used (after classic morphological data) to argue for particular phylogenetic hypotheses. These data include mainly cranial (e.g., **Schoch, 2002, 2006**; **Schoch and Carroll, 2003**; **Schoch and Milner, 2004**; **Anderson, 2007**; **Carroll, 2007**; **Germain and Laurin, 2009**) and autopodial ossification sequences (e.g., **Fröbisch et al., 2007, 2015**). Ossification sequences of other parts of the skeleton, like the vertebrae, shoulder girdle and scales, are also documented in a few Paleozoic stegocephalians (e.g., **Carroll et al., 1999**; **Witzmann, 2006**; **Anderson, 2007**; **Carroll, 2007**; **Olori, 2013**), not to mention finned tetrapodomorphs (**Cloutier, 2010**), but these have played a minor role in the controversy about the origin of extant amphibians. Recently, **Danto et al. (2019)** concluded that vertebral ossification sequences varied too quickly and could not be used to assess the origin of lissamphibians. This study relies on both cranial and appendicular ossification sequences and compares their implications for tetrapod phylogeny.

## Material and methods

### Ossification sequence data

From all the literature we could access, we compiled the most extensive database on ossification sequences for osteichthyans that exists to date. The most useful sources for extant taxa included compilations: **Harrington et al. (2013)** for amphibians, **Weisbecker and Mitgutsch (2010)** for anurans, **Hugi et al. (2012)** for squamates, **Maxwell et al. (2010)** for birds, and **Koyabu et al. (2014)** and **Weisbecker (2011)** for mammals. The cranial and appendicular sequences of Permian temnospondyls (the stereospondylomorphs *Sclerocephalus* and *Archegosaurus*, the non-branchiosaurid “branchiosaur” *Micromelerpeton* and the branchiosaurids “*Melanerpeton*” *humbergense, Apateon caducus* and *A*. *pedestris*) were assembled from several references cited in the **Appendix**; note that the two *Apateon* species are each represented by two different sequences scored after populations from two separate paleo-lakes (Erdesbach and Obermoschel) in which both species occur. Appendicular ossification sequences of the lepospondyls *Microbrachis* and *Hyloplesion* are incorporated from **Olori (2013)**, that for the finned tetrapodomorph *Eusthenopteron* was combined from **Cote et al. (2002)** and **Leblanc and Cloutier (2005)**.

All sources of our sequence data can be found in the **Appendix**. The sequences themselves and the phylogenetic trees corresponding to the tested hypotheses are included in the **Supplementary information**. The sequences were not used to generate the tree topology or the branch lengths (which represent evolutionary time); the tree is compiled from published sources (provided below) which did not use any ossification sequences in their phylogenetic analyses.

The software we used to compute AICc weights, the CoMET module (**Lee et al., 2006**) for Mesquite 3.6 (**Maddison and Maddison, 2018**), cannot handle missing data. This unfortunately meant we had to discard much information. In order to keep as many taxa as possible in the analysis, we first compiled a matrix (not shown) of 244 taxa and 213 characters. All of these characters are positions of skeletal elements (cranial, appendicular, axial and others) in ossification sequences, standardized between 0 and 1 following **Germain and Laurin (2009)**, as explained below. Of these, we kept characters that were scored in the Paleozoic taxa in our initial database, and extant taxa that were scored for the same sets of characters. This resulted in two initial datasets, one of cranial and one of appendicular sequences (it was not possible to include both sets of sequences together because this would have left too few taxa in the matrix).

In the end, however, we were left with three overlapping cranial datasets. The largest cranial dataset we could make, dataset 2 of **Table 1**, has 105 taxa (103 extant, plus the two species of *Apateon* scored from Erdesbach) and seven characters: the appearance times of the premaxilla, maxilla, nasal, parietal, pterygoid, exoccipital and squamosal bones. It lacks *Sclerocephalus*, which cannot be scored for the appearance time of the squamosal. This is unfortunate because *Sclerocephalus* is one of only three extinct taxa for which a usable cranial ossification sequence is known at all, and further because it occupies a special place in the DH2, according to which it lies on the caecilian stem. We attempted to compensate for this deficiency by assembling two more cranial datasets: dataset 1, which contains 107 taxa (104 extant, *Apateon* spp. from Erdesbach, and *Sclerocephalus*) but only six characters by lacking the squamosal, and dataset 5, which includes 84 taxa (81 extant, *Apateon* spp. from Erdesbach, and *Sclerocephalus*) and eight cranial characters (the vomer and the frontal bone are added to the six of dataset 1).

**Table 1.** List of datasets used in this paper. All are subsets of our global compilation that were selected to meet the requirement of the method used (missing data cannot be handled). The temnospondyl species *Apateon caducus* and *A*. *pedestris* are included in all datasets, but scored after populations from two different paleo-lakes in which both species occur.

| Dataset number | 1 | 2 | 3 | 4 | 5 |
| --- | --- | --- | --- | --- | --- |
| Type of characters | cranial | cranial | appendicular | appendicular | cranial |
| Number of characters | 6 | 7 | 7 | 4 | 8 |
| Number of taxa | 107 | 105 | 62 | 65 | 84 |
| *Sclerocephalus* | yes | no | yes | yes | yes |
| Source of data for *Apateon* | Erdesbach | Erdesbach | Obermoschel | Erdesbach and Obermoschel | Erdesbach |
| Additional Paleozoic taxa | None | None | *Archegosaurus*, *Micromelerpeton*, *Hyloplezion*, *Microbrachis*, *Eusthenopteron* | *Archegosaurus*, *Micromelerpeton*, "*Melanerpeton*" *humburgense*, *Hyloplezion*, *Microbrachis*, *Eusthenopteron* | None |
| Tables in which it is used | 2, 5 | 3, 6 | 4, 8 | 4, 9 | 7 |

**Table 2.** Support (AICc and AICc weights) for six evolutionary models given our reference tree (LH) and dataset 1 (see Table 1). Dataset 1 comprises six cranial characters (nasal, parietal, squamosal, maxilla, pterygoid, and exoccipital) scored in 107 taxa, including the temnospondyl *Sclerocephalus*. This was performed on the tree representing the LH (lepospondyl hypothesis), but doing this on other trees leads to similar results. Numbers presented with four significant digits; best values in boldface. “Distance” refers to keeping the original branch length (which represent evolutionary time), “equal” sets all branch lengths (internal and terminal) to 1, “free” infers them from the data. Abbreviations: k, number of estimable parameters; L, likelihood; wi, weight; Δ_*i*_, difference of AICc from that of the Pure-Phylogenetic / Equal model.

| Evolutionary model | AIC | L | k | AICc | $\Delta_i$ AICc | $w_i(\text{AICc})$ |
| --- | --- | --- | --- | --- | --- | --- |
| Pure-Phylogenetic / Distance | -584.4 | 293.2 | **1** | -583.4 | 641.2 | $5.85 E^{-140}$ |
| Pure-Phylogenetic / Equal (speciational) | **-1225.6** | **613.8** | **1** | **-1224.6** | **0** | **1.000** |
| Pure-Phylogenetic / Free | $2.000 E^{10}$ | $-1.000 E^{10}$ | 486 | $2.000 E^{10}$ | $2.000 E^{10}$ | $< E^{-165}$ |
| Non-Phylogenetic / Distance | -473.6 | 237.8 | **1** | -472.6 | 752.0 | $4.97 E^{-164}$ |
| Non-Phylogenetic / Equal | -959.9 | 481.0 | **1** | -958.9 | 265.7 | $2.02 E^{-58}$ |
| Non-Phylogenetic / Free | $2.000 E^{10}$ | $-1.000 E^{10}$ | 244 | $2.000 E^{10}$ | $2.000 E^{10}$ | $< E^{-165}$ |

For the appendicular characters, in addition to dataset 3 which contains seven characters (humerus, radius, ulna, ilium, femur, tibia and fibula) and 62 taxa (54 extant, *Apateon* spp. from Obermoschel, *Sclerocephalus, Archegosaurus, Micromelerpeton, Hyloplesion, Microbrachis* and *Eusthenopteron*), another (dataset 4) includes only four characters (radius, ulna, ilium, and femur), but it features 65 sequences, the additional data being *Apateon* spp. from Erdesbach and “*Melanerpeton*” *humbergense*. See **Table 1** for a list of these datasets and the **Supplementary information** for the datasets themselves.

The data loss in these various datasets is not as severe as it may first seem, because most of the characters that have been excluded from these analyses had less than 10% scored cells (sometimes less than 1%), and most of them could not be scored for any temnospondyl or lepospondyl, so they could not have helped resolve the main question examined in this study.

The order in which the sampled cranial bones ossify varies substantially in our sample of taxa, but based on simple (not phylogenetically-weighted) average position, the frontal appears first, followed closely by the premaxilla, parietal, and maxilla (in close succession), and then by the squamosal, exoccipital, pterygoid, and last by the nasal. However, each of these bones ossifies first (among these bones; not necessarily in the whole skeleton) in at least one of the included taxa. Among the appendicular bones, there is more variability; each ossifies first in at least one of the 62 sampled taxa, and three (radius, ulna and ilium) ossify last in at least one taxon.

Due to the homology problems between the skull bones of tetrapods and actinopterygians and missing data, we had to omit all actinopterygians from our analyses. As cranial ossification sequences remain poorly documented for extant finned sarcopterygians, except perhaps lungfish, whose skull bones seem mostly impossible to homologize (**Criswell, 2015**), our analyses of those data are restricted to limbed vertebrates. However, for appendicular data, we were able to include the Devonian tristichopterid *Eusthenopteron foordi*.

Unfortunately, the only cranial ossification sequence available for any supposed lepospondyl, that of the aïstopod *Phlegethontia longissima*, is documented from only three ossification stages (**Anderson et al., 2003**; **Anderson, 2007**). This poses a problem for our analysis method, which assumes that character evolution can be modeled as Brownian motion; this assumption is decreasingly realistic as the number of character states (sequence positions) decreases, because the resulting distribution deviates increasingly from that of a continuous character. Furthermore, some recent anatomical restudies and phylogenetic analyses suggest that aïstopods are not lepospondyls, but early-branching stem-stegocephalians (**Pardo et al., 2017b**; **Pardo et al., 2018**; **Clack et al., 2019**; **Mann et al., 2019**).

The low taxon sample is more limiting for this analysis than the low character sample. However, as explained below, the absence of lepospondyl sequences in our cranial dataset does not preclude testing the six hypotheses (TH, PH1, PH2, DH1, DH2, LH; see above or **Figure 1** for the explanation of these abbreviations) because each of these six hypotheses makes different predictions about where temnospondyls and caecilians fit relative to other taxa. Thus, in the absence of lepospondyls in our dataset, the tests of these hypotheses are somewhat indirect and inference-based, but they remain possible. Our tests based on appendicular data include two lepospondyls (*Hyloplesion longicostatum* and *Microbrachis pelikani*), but the absence of caecilians in that dataset proves more limiting than the absence of lepospondyls in the cranial dataset because the TH, DH1 and DH2 become indistinguishable (**Fig. 1c, g, h**). However, the presence of the temnospondyl *Micromelerpeton* allows us to test two variants of the TH/DH distinguished by the monophyly (e.g., **Ruta and Coates, 2007**) or polyphyly (e.g., **Schoch, 2019**) of “branchiosaurs” (the temnospondyls *Apateon*, “*Melanerpeton*” *humbergense* and *Micromelerpeton*).

### Sensitivity analysis for sequence polymorphism

Given the potential impact of intraspecific variability in ossification sequence on inferred nodal sequences and heterochrony (**Olori, 2013**; **Sheil et al., 2014**), we compiled two consensus sequences for *Apateon caducus* and *A*. *pedestris* each, representing two localities where both species occur, the paleo-lakes of Erdesbach (**Schoch, 2004**) and Obermoschel (**Werneburg, 2018**). Based on dataset 4 (see **Table 1**), we incorporated these into a global and two separate analyses (one analysis per locality) to determine the impact of the observed variability. As detailed above, incorporating the sequences from Erdesbach reduced the number of characters from seven to only four because the software used cannot handle missing data (see above and below), but this information loss is compensated by the great increase in number of sequences from extinct taxa (eleven instead of two, when counting the sequences of *Apateon* from both localities separately) and the fact that this includes some lepospondyls (see above and below). It would have been even better to perform a sensitivity analysis incorporating variability for all taxa for which such information was available, but given the scope and nature of our study, this would have been exceedingly time-consuming and is best left for the future.

### Standardization of the data

Given that various taxa differ in their numbers of bones and that the resolution of the sequences is also variable between taxa, these data needed to be standardized to make comparisons and computations meaningful, as suggested by **Germain and Laurin (2009)**. Note that we performed this standardization on the complete dataset of characters, before filtering for data completeness. This complete dataset (not shown) includes 213 cranial, appendicular and other characters, but no taxon is scored for all characters, because that matrix has much missing data. For instance, the most completely scored taxon, *Amia calva*, still has 57.4% missing data (more than half), which indicates that 92 characters were scored for this taxon, including several ties (the resolution was 41 positions, so they varied by increments of 0.025 or 2.5% of the recorded ontogeny). We did not re-standardize after filtering characters out because we believe that the initial standardization better reflects the relative position of events in development than a standardization based on only seven events in ontogeny. Because of this, some characters in the reduced datasets lack states 0 or 1 for some taxa. This is simply because the first or last events in the ontogenetic sequence were filtered out. Thus, we used the position in the sequence (from first to last, in the complete dataset) and standardized this relative sequence position between 0 and 1 using the formula given by **Germain and Laurin (2009)**. The standardized sequence position (*X*_*s*_) is:

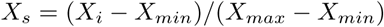

where:

*X*_*i*_ is the position of a given bone in the sequence

*X*_*min*_ is the lowest position in the sequence (generally 0 or 1)

*X*_*max*_ is the highest position in the sequence (for instance, if there are 20 bones, *X*_*min*_ is 1 and the sequence is completely resolved, *X*_*max*_ = 20).

This yields a standardized scale that varies between 0 and 1 for each taxon, in which 0 and 1 are the positions of the first and last events in the sequence, respectively. For instance, for *Ambystoma maculatum* (an extant urodele), in the original dataset, the first events (tied) were the ossification of premaxilla, vomer, dentary and coronoid (standardized position: 0); the last event was the articular (standardized position: 1), and there is a resolution of 12 positions (hence, increments of 0.0909 or 1/11). However, in the final dataset of 7 characters, the articular is absent; hence, the first bone in the sequence is the premaxilla, at a standardized position of 0, and the last is the nasal, as a standardized position of 0.8181 because all events in position 1 (articular) and 0.9091 (stapes) have been filtered out.

We also experimented with using size (skull length) or developmental stage as standards, but this led to lower sequence resolution because body size is not available for all sequence positions and for all taxa (results not shown), so we worked only with sequences standardized by position. Given that our data filtering procedure retains few data (only six, seven or eight characters for the cranial dataset, and four or seven characters for the postcranial dataset), it is important to use the method that discards the least amount of data, and this was achieved by using sequence position. We do not imply that standardizing by size is not recommended in general. On the contrary, if good body size data were available for all taxa and all developmental stages, this should be a better strategy, and only having access to absolute time should be even better. However, practical limitations of data availability prevent us from using these methods now.

Our ossification sequence data (reduced dataset of four to eight characters) of extant and extinct taxa, and the phylogenetic trees we used, are available in the **Supplementary information**.

### Analysis methods

To discriminate between the six hypotheses about the origin of extant amphibians, two methods are available: direct phylogenetic analysis of the sequence data, and comparisons of the tree length (number of steps in regular parsimony, squared length in squared-change parsimony, likelihood, or similar measures) of various trees selected a priori to represent these hypotheses (in these trees, only the position of caecilians and extinct taxa, here temnospondyls and lepospondyls, varies). We used both approaches but expected the second to perform much better because relatively few data are available, and thus, phylogenetic analysis of such data is unlikely to provide a well-resolved tree.

For the first approach, we first transformed the standardized sequence positions back into discrete characters using formulae in a spreadsheet and scaled the characters so that the highest state in all would be 9. This ensures that each character has equal weight in the analysis, regardless of its variability in the ossification sequence. The characters were ordered to reflect the assumed evolutionary model (ontogenetic timing is a quantitative character that was discretized) and because for such characters, ordering yields better results (**Rineau et al., 2015, 2018**; see discussion in **Marjanović and Laurin, 2019**). The resulting data matrices (one for cranial and another for appendicular characters, both with seven characters each) were analysed using parsimony in PAUP* 4.0a165 (**Swofford, 2019**). We used the TBR (tree bisection-reconnection) branch swapping algorithm and performed a search with 50 random addition replicates (or several such searches, for the cranial data) while holding two trees at each step and with a maximum number of trees set at one million. For cranial data, the main search lasted about 100 hours on a MacBook Pro Retina with a 2.5 GHz iCore 7 quadri-core processor and 16 GB RAM. The exact search time cannot be reported because PAUP* crashed after saving the trees to a file for one of the longest runs (several analyses were made, over several days), but before the log could be saved. The analysis of the seven appendicular characters was much faster (27 minutes and a half), presumably because that matrix has fewer taxa (62 instead of 105).

For the second approach (comparison of fit of various trees selected a priori to reflect previously published hypotheses), we used the CoMET module (**Lee et al., 2006**) for Mesquite 3.6 (**Maddison and Maddison, 2018**) to test the relative fit of the data on trees representing the six hypotheses. CoMET calculates the likelihood and the AIC (Akaike Information Criterion) of nine evolutionary models given continuous data and a tree. Note that our data only represent an approximation of continuous data; if standardization had been performed on developmental time or body size, the data would actually have been continuous. Standardization was carried out using sequence position because of data limitation problems, so the data actually follow a decimalized meristic scale. However, the difference between these situations decreases as the number of sequence positions increases, and our global scale includes up to 41 positions (and an average of 10.9 positions), so our data should approximate a continuous distribution sufficiently well for our analyses to be valid. This consideration prevents us from adding the highly apomorphic aïstopod *Phlegethontia*, for which only three cranial ossification stages are known (**Anderson et al., 2003**; **Anderson, 2007**); moreover, five of the eight bones included in our analyses appear in the last two of these stages, one of the relevant bones (vomer) is absent and two (parietal and exoccipital) are not present as separate ossifications, which would create additional missing data. In that case, the very low number of stages would create strong departures from the assumption of continuous data. This would probably create statistical artifacts, and the uncertainty about the position of *Phlegethontia* (**Pardo et al., 2017b**; **Pardo et al., 2018**; **Clack et al., 2019**; **Marjanović and Laurin, 2019**) would complicate interpretation of the results.

The nine models evaluated by CoMET are obtained by modifying the branch lengths of the reference tree. Thus, branches can be set to 0 (for internal branches only, to yield a non-phylogenetic model), to 1 (equal or speciational model), left unchanged from their original length (gradual evolution in our case, where the original lengths represent geologic time), or set free and evaluated from the data (free model). This can be applied to internal and/or external branches, and various combinations of these yield nine models (**Lee et al., 2006, fig. 1**). Among these nine models two have been frequently discussed in the literature and are especially relevant. The first is gradual evolution, in which branch lengths (here representing evolutionary time) have not been changed. The second is the speciational model, in which all branches are set to the same length because changes are thought to occur at speciation events, which are typically equated with cladogeneses in evolutionary models (**Bokma et al., 2016**). This model has some similarities with Eldredge and Gould’s (1972) punctuated equilibria (though a model with one internal branch stemming from each node set to 0 and the other set to 1 would be even closer to the original formulation of that model). In this study, we assessed the fit of six of the nine models covered by CoMET; the other three (the punctuated versions of distance [original branch length], equal and free), in which one of each pair of daughter-lineages has a branch length of zero, could not be assessed due to problems in the current version of CoMET and possibly the size of our dataset.

Provided that the same evolutionary model is optimal for all compared phylogenetic hypotheses (this condition is met, as shown below), the AIC weights of the various trees under that model can be used to assess the support for each tree. In such comparisons, the topology is part of the evolutionary model, and the data are the sequences. These comparisons can show not only which tree is best supported, but how many times more probable the best tree is compared to the alternatives. This quantification is another reason to prefer this approach over a phylogenetic analysis (performed below, but with the poor results that we anticipated), which can at best yield a set of trees showing where the extinct taxa most parsimoniously fit (if we had dozens of characters, this might be feasible). Comparisons with other hypotheses through direct phylogenetic analysis are not possible. Given the small sample size (which here is the number of characters), we computed the corrected AIC (AICc) and the AICc weights using the formulae given by **Anderson and Burnham (2002)** and **Wagenmakers and Farrell (2004)**.

Our tests make sense only in the presence of a phylogenetic signal in the data. In addition to the test of evolutionary model in CoMET mentioned above (which tests non-phylogenetic as well as phylogenetic models), we performed a test based on squared-change parsimony (**Maddison, 1991**) and random taxon reshuffling (**Laurin, 2014**). For this test, we compared the length of the LH (lepospondyl hypothesis; **Fig. 1d**) reference tree (with and without *Sclerocephalus*) to a population of 10,000 random trees produced by taxon reshuffling.

It could be argued that using other methods (in addition to the method outlined above) would have facilitated comparisons with previous studies. However, the two main alternative methods, event-pair cracking with Parsimov (**Jeffery et al., 2005**) and Parsimov-based genetic inference (PGI; **Harrison and Larsson, 2008**), have drawbacks that led us to not using them. Our objections against event-pair cracking with Parsimov were detailed by **Germain and Laurin (2009)**. In short, that method requires an unnecessary decomposition of sequences into event pairs, and it cannot incorporate absolute timing information (in the form of time, developmental stage or body size, for instance) or branch length information. More importantly, the simulations performed by **Germain and Laurin (2009)** showed that event-pair cracking with Parsimov yields more artefactual change and has lower power to detect real sequence shifts. That method is also problematic when trying to infer ancestral sequences and can lead to impossible ancestral reconstructions (e.g. A occurs before B, B occurs before C, and C occurs before A), as had been documented previously (**Schulmeister and Wheeler, 2004, p. 55**). This would create problems when trying to compare the fit of the data on various phylogenetic hypotheses. The performance of Parsimov-based genetic inference (PGI; **Harrison and Larsson, 2008**) has not been assessed by simulations, but it rests on an edit cost function that is contrary to our working hypothesis (that the timing of developmental events can be modeled with a bounded Brownian motion model, which is assumed by continuous analysis). More specifically, **Harrison and Larsson (2008, p. 380)** stated that their function attempts to minimize the number of sequence changes, regardless of the magnitude of these changes. We believe that disregarding the size of changes is unrealistic, as shown by the fact that Poe’s (2006) analyses of thirteen empirical datasets rejected that model (which he called UC, for unconstrained change) in favor of the model we accept (A J for adjacent states, which favors small changes over large ones). Furthermore, analyses of ossification sequence data using techniques for continuous data as done here (see above) have been performed by an increasingly large number of studies (e.g., **Skawinski and Borczyk, 2017**; **Spiekman and Werneburg, 2017**; **Werneburg and Geiger, 2017**; just to mention papers published in 2017), so the issue of ease of comparisons of our results with other studies is not as serious as it would have been only a few years ago, and it should be decreasingly so in the future.

### Reference phylogenies

We built a reference timetree that attempts to capture the established consensus (**Fig. 2**; see the next paragraphs for the sources). The tree was compiled in Mesquite versions up to 3.6 (**Maddison and Maddison, 2018**) and time-calibrated using the Stratigraphic Tools module for Mesquite (**Josse et al., 2006**). For consistency and to avoid the effects of gaps in the fossil record, we used molecular divergence dates whenever possible. The tree had to be time-scaled because many of the evolutionary models that we fit on the tree in the first series of tests (to determine which evolutionary model can be used to compare the fit of the hypotheses) use branch lengths to assess model fit. Note that our procedure requires estimating divergence times between all taxa (geological ages of all nodes). When taxa are pruned, branch lengths are adjusted automatically. The main sources we used for topology and divergence times (and hence branch lengths) are as follows.

The phylogeny of lissamphibians follows the work of **Jetz and Pyron (2018)**. However, several other sources have been used for the temporal calibration of the tree: **Germain and Laurin (2009)** was used for the urodeles, whereas **Feng et al. (2017)**, supplemented by **Bossuyt and Roelants (2009)** and **Pyron (2014)**, was used for the anurans as well as more rootward nodes (Batrachia, Lissamphibia, Tetrapoda; also Amniota). **Marjanović and Laurin (2014)** was used for the Ranidae, Ceratophryidae and Hylidae.

The sediments that have preserved the temnospondyls *Apateon* and *Sclerocephalus* are not easy to correlate with each other or with the global chronostratigraphic scale. Combining stratigraphic information from **Schoch (2014b), Schneider et al. (2015)** and **Werneburg (2018)**, we have placed all three sampled species (*A*. *pedestris, A*. *caducus, S*. *haeuseri*) at the Sakmarian/Artinskian stage boundary (Permian; 290.1 Ma ago); combining stratigraphic information from **Schneider et al. (2015)** with the phylogeny in **Schoch (2014b)**, we have tentatively placed the divergence between the two *Apateon* species (which are not sister-groups: **Schoch, 2014b**) at the Kasimovian/Gzhelian stage boundary (Carboniferous; 303.7 Ma ago). The age of the last common ancestor of *Apateon* and *Sclerocephalus* depends strongly on temnospondyl phylogeny, which remains unresolved (**Pardo et al., 2017a**; **Marjanović and Laurin, 2019**; and numerous references in both); as a compromise between the various options, we have provisionally placed it at the boundary between the Early and the Late Carboniferous (Serpukhovian/ Bashkirian, 323.2 Ma ago) where applicable.

We sampled many extant amniotes to achieve broad coverage of Tetrapoda. For the birds, **Pons et al. (2005)** was used for the Laridae, **Wang et al. (2013)** for the Phasianidae and **Gonzalez et al. (2009)** for the Anatidae. The temporal calibration was taken from **Prum et al. (2015)** as recommended by **Berv and Field (2018)**; gaps were filled in using the database www.birdtree.org.

Several papers, mainly **Tarver et al. (2016)**, were used for the phylogeny and divergence times of mammals. For the Muridae, three references were used: **Lecompte et al. (2008), Zhuang et al. (2015)**, and **Lu et al. (2017)** for the position of two taxa: *Mesocricetus auratus* and *Peromyscus melanophrys*. Other species were placed following the work of **Meredith et al. (2011)**, which also gives divergence times. We caution, however, that all available molecular dates for Paleogene and earlier mammal nodes are controversial and may be overestimates (**Berv and Field, 2018**; **Phillips and Fruciano, 2018**).

Three references were also used to integrate squamates in the phylogenetic tree and for the calibration of divergence times: **Brandley et al. (2005), Rabosky et al. (2014), Reeder (2003). Sterli et al. (2013)** was used for turtles.

For turtles, there is now a near-consensus that they are diapsids, a hypothesis that is not necessarily incompatible with an origin among “parareptiles” (**Laurin and Piñeiro, 2017**). Thus, following most recent phylogenetic analyses of molecular data (e.g., **Hugall et al., 2007**; **Irisarri et al., 2017**), we have inserted them as the sister-group of Archosauria.

We disagree with several of the calibration dates in **Irisarri et al. (2017)**, which often appear unreasonably old. For instance, they place the divergence between caecilians and batrachians and the divergence between anurans and urodeles in the Early Carboniferous, around 330 and 320 Ma, respectively, but our thorough analyses of the fossil record, with due consideration of its incompleteness, suggest significantly more recent dates, in the Permian (**Marjanović and Laurin, 2007, 2008, 2014**). This is not surprising because some of the dating constraints used by **Irisarri et al. (2017, table S8)** are wrong. For instance, they enforced a minimal divergence age between cryptodiran and pleurodiran turtles of 210 Ma (Late Triassic), but all analyses of the last fifteen years (e.g., **Sterli et al., 2013, 2018**) strongly suggest that the oldest known turtles that fit within this dichotomy date from the Late Jurassic, less than 165 Ma. The divergence between humans and armadillos (boreotherian and xenarthran placentals) was constrained to the middle of the Cretaceous (95.3–113 Ma), based on outdated literature that assigned a wide variety of stem-eutherians to highly nested positions in the placental crown; there are currently no clear placentals known from any Cretaceous sediments even as young as 66 Ma (e.g., **Halliday et al., 2016, 2017**; **Davies et al., 2017**; **Phillips and Fruciano, 2018**), barely half the age of the older end of the constraint range. Conversely, the divergence between diapsids (hence sauropsids) and synapsids had a minimal age constraint of 288 Ma (Early Permian), which is much too young given the presence of sauropsids (and presumed synapsids) in Joggins, in sediments that have recently been dated (**Carpenter et al., 2015**) around 317–319 Ma (early Late Carboniferous). Thus, we have not used divergence dates from that source.

To discriminate among the hypotheses on lissamphibian origins, we inserted the temnospondyl *Apateon* in the tree where each predicts that it should be (**Fig. 1c–h**). Thus, according to the TH (temnospondyl hypothesis; **Fig. 1c**), *Apateon* lies on the lissamphibian stem. Under the LH (lepospondyl hypothesis; **Fig. 1d**), *Apateon* lies on the tetrapod stem. Under both versions of the DH (diphyly hypothesis; **Fig. 1g, h**), *Apateon* lies on the batrachian stem. Under both versions of the PH (polyphyly hypothesis; **Fig. 1e, f**), *Apateon* lies on the caudate stem. Within the DH and the PH, both versions of each differ in the position of Gymnophiona. Thus, despite the absence of any lepospondyl in our cranial ossification sequence datasets, our taxonomic sample allows us to test all these competing hypotheses. The appendicular datasets allow more direct tests of some of these hypotheses because they include two lepospondyl taxa, which were likewise placed in trees representing the tested hypotheses (**Fig. 1**).

*Sclerocephalus* is the sister-group of *Apateon* under the LH (**Fig. 1d**), immediately rootward of it (on the lissamphibian stem) under the TH (**Fig. 1c**) and likewise (but on the batrachian stem) under the DH1 (**Fig. 1g**), on the caecilian stem under the DH2 (**Fig. 1h**) and the sister-group of Batrachia (including *Apateon*) under both versions of the PH (**Fig. 1e, f**).

“*Melanerpeton*” *humbergense* (appendicular data only) is the sister-group of *Apateon* in all trees, except under the hypothesis of branchiosaur paraphyly; *Eusthenopteron* (appendicular data only) forms the outgroup in all trees.

The lepospondyls *Microbrachis* and *Hyloplesion*, from both of which only appendicular data are available, form an exclusive clade (**Clack et al., 2019**; **Marjanović and Laurin, 2019**). This clade is the sister-group of Lissamphibia (represented only by Batrachia because caecilians are lacking from the appendicular datasets) under the LH, of Amniota under the TH and both versions of the DH (these three cannot be distinguished due to the absence of caecilians) as well as under the PH1, and of Temnospondyli (including Batrachia) under the PH2 (see the legend of **Figure 1** for an explanation of these abbreviations).

The temnospondyl *Micromelerpeton*, from which likewise only appendicular data are available, forms the sister-group of *Apateon* under the LH. The uncertainty over its phylogenetic position within Dissorophoidea (as the sister-group to the rest, including anurans and urodeles: e.g. **Schoch, 2019**; as the sister-group of *Apateon* + “*Melanerpeton*” *humbergense*: e.g. **Ruta and Coates, 2007**; **Marjanović and Laurin, 2019**) generates two versions of the TH/DH1/DH2 tree for the appendicular dataset. We tested both of these versions against that dataset, for a total of five trees.

To ensure that our analyses were not biased in favor of a given hypothesis, and in case that a continuous evolutionary model were favored, we initially adjusted the branch lengths such that the sum of branch lengths was equal between the compared topologies and that the root was approximately at the same age (in this case in the Tournaisian, the first stage of the Carboniferous). This was done for the trees used to compare the hypotheses using the cranial dataset because if a model incorporating (variable) branch length information had been selected, and if the trees representing the various hypotheses had not all had the same total length (the sum of all branch lengths), the resulting distortions in branch lengths created around the extinct taxa (whose height compared to extant taxa is specified by their geological age) would have introduced another variable influencing the AICc. But given that the selected model ignores branch lengths, this precaution turned out to be superfluous. We have therefore not made these time-consuming adjustments to the additional trees we generated later to analyze the appendicular data.

## Results

In the phylogenetic analysis of cranial data, a single tree island of 22,077 trees of 438 steps was found, only once, so there might be more trees of that length and perhaps even shorter trees. Initially, an island of 22,075 trees was found; we swapped on each of these in a subsequent run, which only recovered two additional trees. Given that slightly longer trees did not differ much from those that we obtained, the low quality of the results (poor congruence with the established consensus about the monophyly of major clades such as squamates, birds, mammals and turtles) and the fact that about four full days of computer time had been spent on analysis of the cranial data, we did not pursue that search further. As expected, the strict consensus tree is poorly resolved (**Fig. 3**). The majority-rule consensus (not shown, but available in the **Supplementary information**) is more resolved but not necessarily better because much of the additional resolution contradicts the established consensus. For the appendicular matrix, 22,757 trees of 164 steps were found. Their strict consensus (**Fig. 4**) deviates even more from the established consensus than the tree obtained from cranial data.

**Figure 3.**
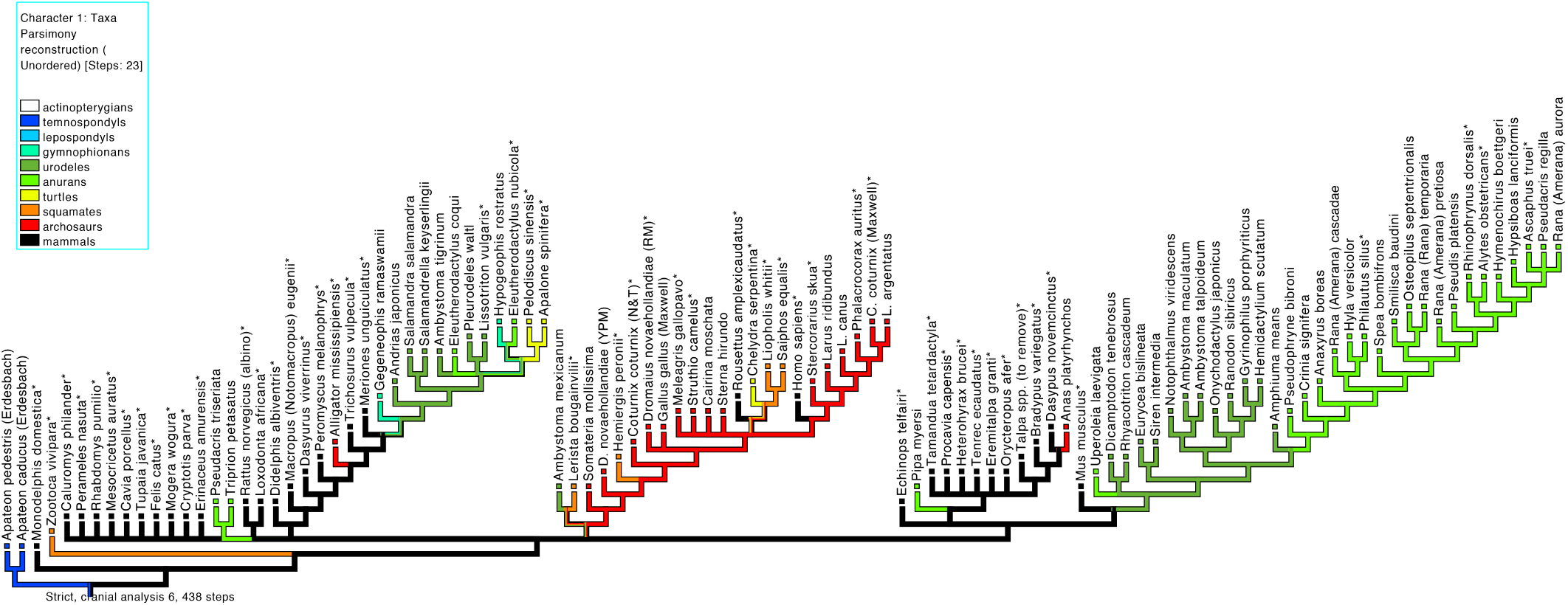
Strict consensus of the most parsimonious trees obtained by analyzing cranial dataset 2. Dataset 2 comprises 105 taxa and seven characters (see **Table 1**). Note that several higher taxa whose monophyly is well-established are paraor polyphyletic here. Abbreviations: C., *Coturnix*; L., *Larus*. Asterisks meaningless.

**Figure 4.**
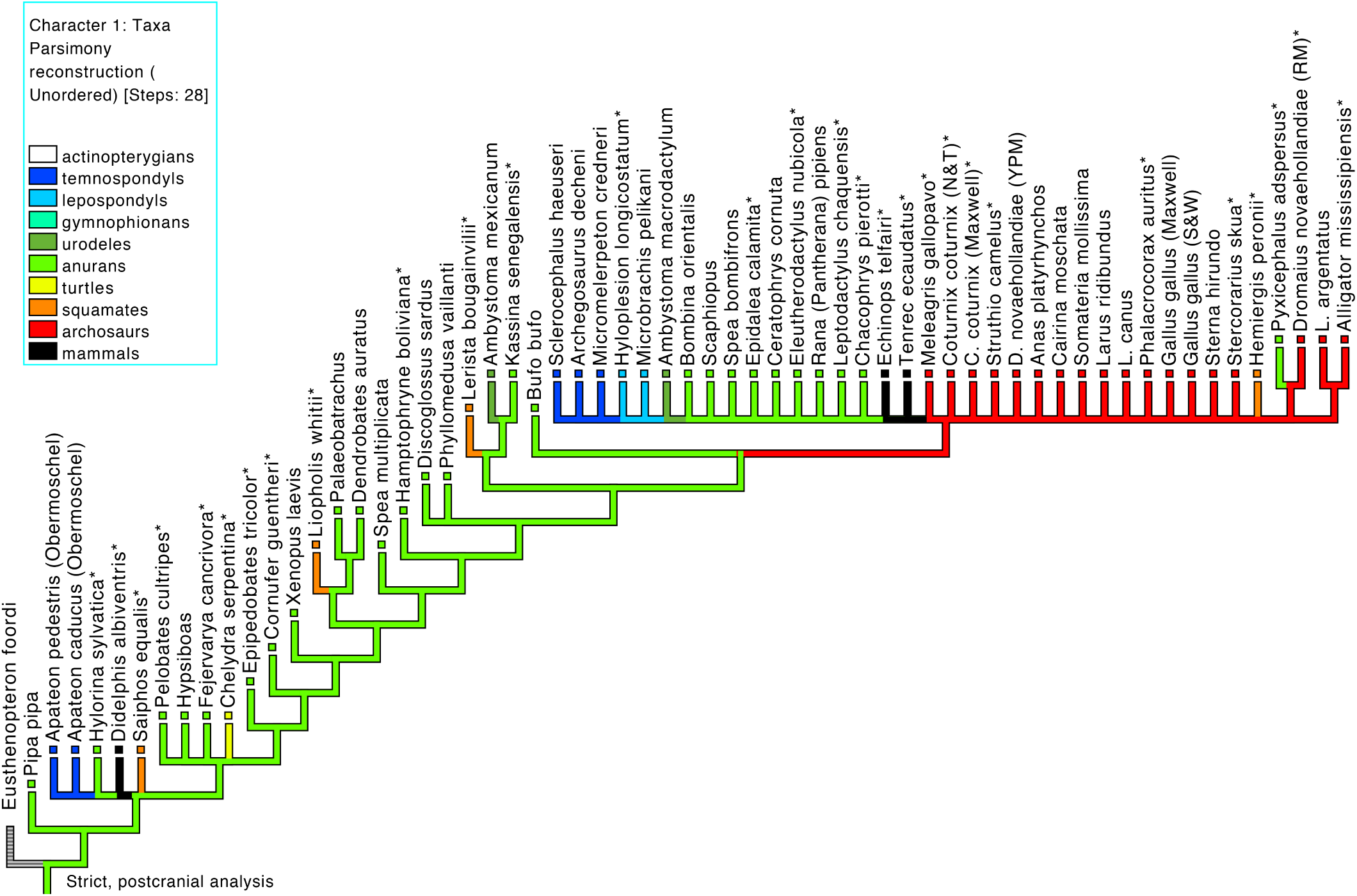
Strict consensus of the most parsimonious trees obtained by analyzing appendicular dataset 3. Dataset 3 comprises 62 taxa and seven characters (see **Table 1**). The phylogenetic signal in these data seems to be lower than in the cranial data. See **Fig. 3** for abbreviations.

This visual assessment of phylogenetic signal through an examination of the consensus trees (**Figs. 3, 4**) is congruent with the test based on squared-change parsimony and random taxon reshuffling (**Laurin, 2004**). Indeed, the latter indicates that the phylogenetic signal in the cranial data is fairly strong, with a probability of less than 0.0001 that the observed covariation between the data and the tree reflects a random distribution (none of the 10,000 random trees generated were as short as the reference tree). However, it is weaker, with a probability of 0.0017, for the appendicular data.

The speciational model of evolution, in which all branch lengths are equal, has overwhelming support among cranial data, whether or not the Permian temnospondyl *Sclerocephalus* (**Table 2**) or the squamosal (**Table 3**) are included (including *Sclerocephalus* adds a second temnospondyl genus, but given that the timing of ossification of the squamosal is unknown in *Sclerocephalus*, including it requires excluding the squamosal from the analysis as described in the Methods section); the five other examined models have AICc weights < 10^−11^. For the appendicular data, the speciational model also has the most support, but that support is not as strong and varies depending on which dataset is analyzed (seven characters or four) and under which phylogenetic hypothesis. In three of the four tests performed, support for the second-best model, the non-phylogenetic/equal model, varied between 5% and 19% (**Table 4**).

**Table 3.**
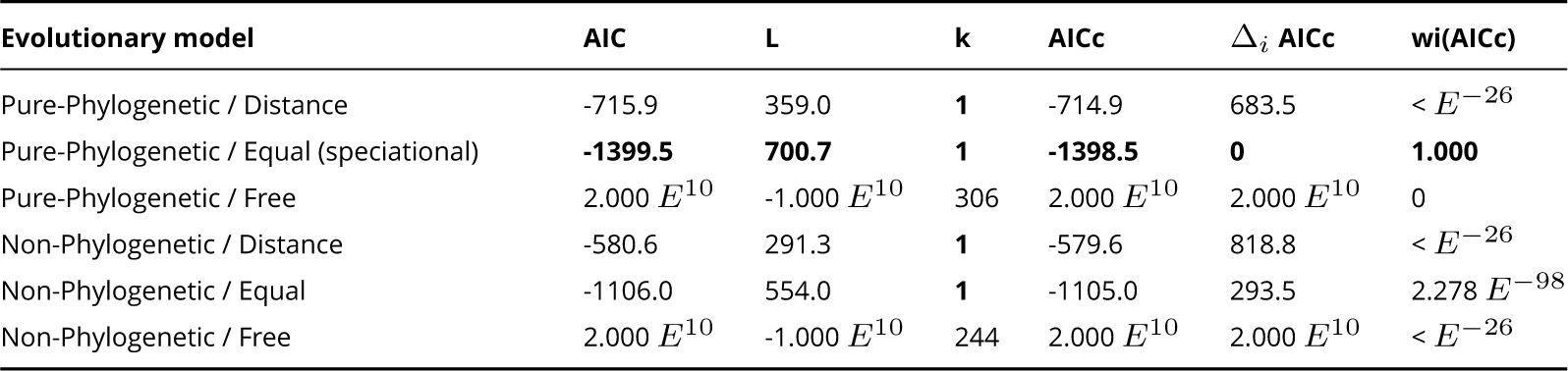
Support (AICc and AICc weights) for six evolutionary models given our reference tree (LH) and dataset 2 (see Table 1). Dataset 2 comprises seven cranial characters (nasal, parietal, squamosal, premaxilla, maxilla, pterygoid, and exoccipital) and 105 taxa, excluding *Sclerocephalus*. Abbreviations and boldface as in **Table 2**.

**Table 4.** AICc weights showing relative support for six evolutionary models given our appendicular datasets (3 and 4; see Table 1) and various hypotheses. Because of the number of analyses presented below, only the AICc weights are presented (best values in boldface). Abbreviations: DH, diphyly hypothesis (both versions); LH, lepospondyl hypothesis; TH, temnospondyl hypothesis.

| Evolutionary model | 7 characters, LH | 7 characters, LH | 4 characters, LH | 4 characters, TH/DH |
| --- | --- | --- | --- | --- |
| Pure-Phylogenetic / Distance | 5.1857 $E^{-149}$ | 2.340 $E^{-70}$ | 1.227 $E^{-52}$ | 2.646 $E^{-52}$ |
| Pure-Phylogenetic / Equal | **1** | **0.9335** | **0.94459** | **0.8139** |
| Pure-Phylogenetic / Free | < $E^{-179}$ | 1.598 $E^{-277}$ | 4.012 $E^{-158}$ | 3.002 $E^{-155}$ |
| Non-Phylogenetic / Distance | 7.515 $E^{-179}$ | 4.843 $E^{-52}$ | 2.162 $E^{-42}$ | 7.262 $E^{-42}$ |
| Non-Phylogenetic / Equal | 2.14914 $E^{-64}$ | 6.648 $E^{-2}$ | 5.541 $E^{-2}$ | 0.1861 |
| Non-Phylogenetic / Free | < $E^{-179}$ | < $E^{-179}$ | < $E^{-179}$ | < $E^{-179}$ |

Two main conclusions can be drawn from these tests (**Tables 2–4**). First, given that both of the best-supported models imply equal branch lengths, actual time represented by branches can be ignored, so we compare support of the six competing topologies using only the best-supported model (speciational). This simplifies the discussion, because it means that the original branch lengths are irrelevant (under that model, all branch lengths are equal); unfortunately, the branch length (evolutionary time) data were needed to reach this conclusion. Thus, the only remaining variable is the topology. Second, model fitting, along with the test based on squared-change parsimony and random taxon reshuffling, indicates that the phylogenetic signal in the cranial data is strong, but that it is noticeably weaker in the appendicular data (this is shown mostly by the non-negligible support for the non-phylogenetic/equal model). Thus, comparisons of the fit of the various phylogenetic hypotheses for the cranial data should be more reliable than for the appendicular data. However, given that for several Paleozoic taxa (most importantly both of the sampled lepospondyls), comparisons can be performed only for the appendicular data, these were performed as well.

Using the speciational model, the AICc weights of the six compared topologies indicate that there is strong support in the cranial data for the LH (lepospondyl hypothesis), with an AICc weight of 0.9885 when *Sclerocephalus* is included (**Table 5**) and 0.8848 when the squamosal is included instead (**Table 6**). Of the other topologies, the TH (temnospondyl hypothesis) was by far the best supported, with an AICc weight of 0.01144 (with *Sclerocephalus*) or 0.1056 (with the squamosal), which is 86.44 or 8.38 times less than for the LH. Both versions of the DH (diphyly hypothesis) and of the PH (polyphyly hypothesis) have negligible support (AICc weights < 0.01 when the squamosal is included, < 0.0001 when *Sclerocephalus* is included). The least support is found for the PH2 when *Sclerocephalus* is included, and for the DH1 when the squamosal is included. In both cases, the recently proposed DH2 (**Pardo et al., 2017a**) fares second-worst by a small margin. Notably, the DH1 contradicts the modern consensus on lissamphibian monophyly (**Fig. 1g**), while the PH2 and the DH2 fulfill this constraint from the molecular but not the paleontological point of view, having lissamphibian monophyly with respect to amniotes but not with respect to temnospondyls (**Fig. 1f, h**).

**Table 5.**
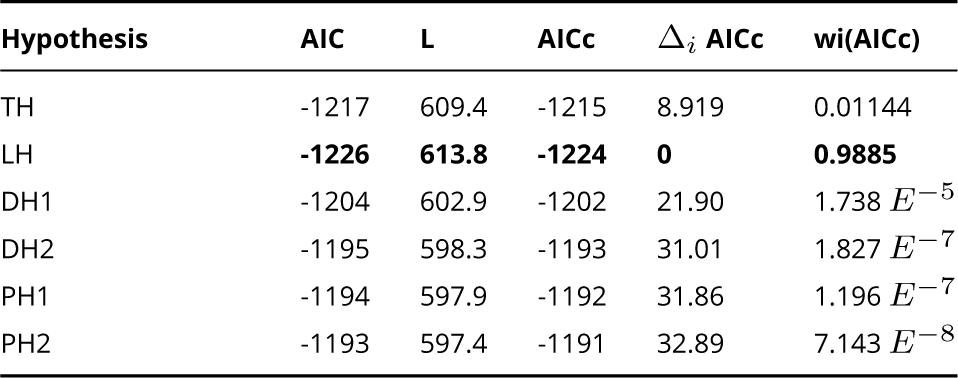
Support (AIC and AICc weights) for the six topologies, reflecting the six hypotheses about the origin of extant amphibians, under the speciational model (called Pure-Phylogenetic / Equal in Tables 2–4), with dataset 1 (see Table 1). Dataset 1 includes six cranial characters (nasal, parietal, squamosal, maxilla, pterygoid, and exoccipital) and 107 taxa (including, among Paleozoic taxa, *Apateon* and *Sclerocephalus*). Abbreviations and boldface as in **Table 2**, except Δ_*i*_: difference of AICc from that of the LH. Hypotheses from top to bottom: TH: monophyletic origin among temnospondyls; LH: monophyletic origin from lepospondyls; DH1: diphyletic origin, caecilians from lepospondyls and batrachians from temnospondyls, as in **Anderson et al. (2008)**; DH2: diphyletic origin, (batrachians and caecilians from different temnospondyls: **Pardo et al., 2017a**); PH1: triphyletic (polyphyletic) origin with anurans and urodeles from different temnospondyls, caecilians from lepospondyls, and lepospondyls closer to Amniota than to Batrachia (**Fröbisch et al., 2007**); PH2: triphyletic (polyphyletic) origin as above, but with lepospondyls and caecilians closer to temnospondyls than to amniotes (**Milner, 1993**), reflecting the well-established lissamphibian monophyly among extant taxa (e.g., **Feng et al., 2017**; **Irisarri et al., 2017**).

**Table 6.** Support (AIC and AICc weights) for the six topologies, reflecting the six hypotheses about the origin of extant amphibians, for dataset 2 (see Table 1). Dataset 2 includes seven cranial characters (nasal, parietal, squamosal, premaxilla, maxilla, pterygoid, and exoccipital) and 105 taxa, excluding *Sclerocephalus* (among Paleozoic taxa, only *Apateon* is present). Abbreviations, boldface and hypotheses as in **Tables 2** and **5**.

| Hypothesis | AIC | L | AICc | $\Delta_i$ AICc | wi(AICc) |
| --- | --- | --- | --- | --- | --- |
| TH | -1395 | 698.6 | -1394 | 4.251 | 0.1056 |
| LH | **-1399** | **700.7** | **-1398** | **0** | **0.8848** |
| DH1 | -1384 | 693.1 | -1383 | 15.203 | $4.42 E^{-4}$ |
| DH2 | -1385 | 693.6 | -1384 | 14.315 | $6.89 E^{-4}$ |
| PH1 | -1387 | 694.5 | -1386 | 12.404 | $1.792 E^{-3}$ |
| PH2 | -1390 | 695.8 | -1388 | 9.792 | $6.615 E^{-3}$ |

A slightly different dataset (only 84 taxa, but eight cranial characters – excluding the squamosal but including the frontal and the vomer – and *Apateon* sequences for both species from Erdesbach rather than Obermoschel) provides even stronger support for the LH, with an AICc weight of 0.9935 (**Table 7**). The next best-supported topology, which simultaneously represents the TH, DH1 and DH2 (due to the absence of caecilians from this dataset), has an AICc weight of only 0.0065.

**Table 7.** Support for the various hypotheses about amphibian origins for dataset 5 (see Table 1). Dataset 5 includes eight cranial characters (frontal added) and 84 taxa, with *Apateon* sequences from Erdesbach (in addition to *Sclerocephalus* among Paleozoic taxa). Abbreviations, boldface and hypotheses as in **Tables 2** and **5**. Because of the taxon sample, only three topologies can be tested.

| Hypothesis | AIC | L | AICc | $\Delta_i$ AICc | wi(AICc) |
| --- | --- | --- | --- | --- | --- |
| LH | **-1296** | **649.0** | **-1294** | **0** | **0.9935** |
| TH, DH1, DH2 | -1286 | 644.0 | -1284 | 10.061 | $6.493 E^{-3}$ |
| PH1, PH2 | -1274 | 638.0 | -1272 | 22.038 | $1.628 E^{-5}$ |

The appendicular data are available in far more Paleozoic taxa than the cranial data; these include *Sclerocephalus haeuseri, Archegosaurus decheni*, and the non-branchiosaurid “branchiosaur” *Micromelerpeton credneri* among temnospondyls, the lepospondyls *Hyloplesion longicaudatum* and *Microbrachis pelikani*, and the tristichopterid finned stem-tetrapodomorph *Eusthenopteron foordi*, in addition to the same two species of *Apateon* as for the cranial datasets, *A*. *caducus* and *A*. *pedestris*. Analysis of these data (seven characters: humerus, radius, ulna, ilium, femur, tibia and fibula) yields surprising results, with the PH2 having the most support, with an AICc weight of 0.7978 when using the dataset of seven bones (**Table 8**). The TH, DH1 and DH2 with “branchiosaur” monophyly are collectively (they cannot be distinguished with that taxonomic sample) the second-best hypotheses with that dataset, with an AICc weight of only 0.1874. The least-supported hypothesis with these data is the TH/DH with “branchiosaur” polyphyly.

**Table 8.** Support (AICc weights) for the various hypotheses about amphibian origins according to dataset 3 (see Table 1). Dataset 3 features seven appendicular characters (humerus, radius, ulna, ilium, femur, tibia and fibula) and 62 taxa, including several Paleozoic taxa (the temnospondyls *Archegosaurus decheni* and *Micromelerpeton credneri*, the lepospondyls *Hyloplesion longicaudatum* and *Microbrachis pelikani*, and the tristichopterid *Eusthenopteron foordi*) in addition to *Apateon* (two species, *A*. *caducus* and *A*. *pedestris*) and *Sclerocephalus haeuseri*. The *Apateon* sequences come from Obermoschel. Abbreviations, boldface and hypotheses as in **Table 5**, except that the TH and both variant of the DH become indistinguishable, but the phylogenetic position of the “branchiosaur” *Micromelerpeton* can be tested.

| Hypothesis | AIC | L | AICc | $\Delta_i$ AICc | wi(AICc) |
| --- | --- | --- | --- | --- | --- |
| LH | -885.0 | 443.5 | -884.2 | 11.808 | $2.177 E^{-3}$ |
| TH, DH (branchiosaur monophyly) | -881.1 | 441.6 | -880.3 | 2.897 | 0.1874 |
| TH, DH (branchiosaur polyphyly) | -886.4 | 444.2 | -885.6 | 15.754 | $3.027 E^{-4}$ |
| PH1 | -888.5 | 445.3 | -887.7 | 8.341 | 0.01232 |
| PH2 | **-896.9** | **449.4** | **-896.1** | **0.000** | **0.7978** |

Using the other postcranial dataset with only four bones (radius, ulna, ilium, and femur) but with more taxa (notably the branchiosaurid temnospondyl “*Melanerpeton*” *humbergense*) shows that intraspecific variation in the postcranial ossification sequences of *Apateon* do not significantly impact our assessment of the support for various hypotheses. Whether both sequences of *Apateon* (from the Erdesbach and Obermoschel localities, which represent separate paleo-lakes) are included (treated as if they were distinct taxa, such as subspecies), or whether either one of these is used in isolation, the PH2 retains the highest support, with AICc weights of 0.62 to 0.65. The LH is a distant second, at 0.20–0.23, but still well ahead of the TH/DH and the PH1, which all receive AICc weights between 0.03 and 0.06 (**Table 9**).

**Table 9.** Effect of the intraspecific variability in ossification sequences of *Apateon* on the support (AICc weight; best values in boldface) for the various hypotheses about amphibian origins. The dataset (number 4; **Table 1**) includes only four appendicular bones (radius, ulna, ilium, and femur) and 63 to 65 taxa but it allows testing the impact of intraspecific variability in ossification sequences in *Apateon*, which are documented in two localities (Erdesbach and Obermoschel). Because of the number of tests presented (15: five topologies x three sets of sequences), only the AICc weights are given. In all tests, the following Paleozoic taxa are present: *Sclerocephalus haeuseri, Archegosaurus decheni*, “*Melanerpeton*” *humbergense, Micromelerpeton credneri, Apateon* (two species, *A*. *caducus* and *A*. *pedestris*) among temnospondyls, *Hyloplesion longicaudatum* and *Microbrachis pelikani* among lepospondyls, and the tristichopterid *Eusthenopteron foordi*. For abbreviations of the hypotheses, see **Table 5**.

| Hypothesis | Erdesbach and Obermoschel | Erdesbach | Obermoschel |
| --- | --- | --- | --- |
| LH | 0.21407 | 0.20169 | 0.22657 |
| TH, DH (branchiosaur monophyly) | 0.05492 | 0.05265 | 0.05532 |
| TH, DH (branchiosaur polyphyly) | 0.03713 | 0.04285 | 0.03342 |
| PH1 | 0.05653 | 0.05491 | 0.05638 |
| PH2 | **0.63735** | **0.64790** | **0.62832** |

## Discussion

### Phylogenetic signal

In his discussion of previous phylogenetic conclusions from ossification sequences (e.g., **Schoch and Carroll, 2003**), **Anderson (2007)** noted that ossification sequences seemed to abound in symplesiomorphies and in autapomorphies of terminal taxa, while potential synapomorphies were scarce. This pessimism was seemingly confirmed by **Schoch (2006)** in a paper that was published after Anderson’s **(2007)** book chapter had gone to press: not only were many similarities in the cranial ossification sequences across Osteichthyes found to be symplesiomorphies, but a phylogenetic analysis of cranial ossification sequences did not recover Mammalia, Sauropsida, Amniota or Lissamphibia as monophyletic. Along with these results, **Schoch (2006)** dismissed another: the position of the temnospondyl *Apateon caducus* (the only included extinct taxon) outside the tetrapod crown-group, i.e. the lepospondyl hypothesis on lissamphibian origins (LH).

While ossification sequences alone may not provide enough data for a phylogenetic analysis, as shown by our results (**Figs. 3, 4**), there is clearly a phylogenetic signal because the taxa are not randomly scattered over the tree. Specifically, our datasets (with much larger taxon samples than in **Schoch, 2006**) fit some tree topologies much better than others. Both the tests using CoMET and squared-change parsimony with random taxon reshuffling overwhelmingly support the presence of a strong phylogenetic signal in the cranial data; the null hypothesis of the absence of a phylogenetic signal can be rejected in both cases, given that it has a probability of < 10^−97^ for the cranial and < 10^−4^ for the appendicular dataset. We conclude that the cranial dataset contains a strong phylogenetic signal, and are therefore cautiously optimistic about future contributions of ossification sequences to phylogenetics. We are less optimistic about the appendicular sequence data, which both tests suggest contains less phylogenetic signal.

The sizable effect on nodal estimates and inferred heterochronies of intraspecific variation found by **Sheil et al. (2014)** in lissamphibians could raise doubts about the robustness of our findings. We have been able to incorporate infraspecific variability in only two terminal taxa (*Apateon caducus* and *A*. *pedestris*), but *Apateon* has played a prominent role in discussions about the significance of cranial ossification sequences on the origins of extant amphibians (**Schoch and Carroll, 2003**; **Schoch, 2006**; **Germain and Laurin, 2009**). Thus, incorporation of intraspecific variability in *Apateon* is presumably much more important than in extant taxa, even though variability in the latter would obviously add to the analysis and should be tackled in the future. The variability in *Apateon* should be exempt from two sources of artefactual variability in ossification sequences discussed by **Sheil et al. (2014)**, namely the way in which the specimens were collected (there can be no lab-raised specimens in long-extinct taxa) and the fixing method used (in this case, fossilization under quite consistent taphonomic conditions). The finding that the results are very similar no matter whether we used the *Apateon* sequences from Erdesbach, Obermoschel, or both, we find very similar results (**Table 9**), is reassuring. In this case, intraspecific variation has negligible impact. However, future studies should attempt to assess the effect of more generalized incorporation of intraspecific variability (in a greater proportion of the OTUs).

Of course, these results do not preclude functional or developmental constraints from applying to the same data. This phenomenon has been documented, among other taxa, in urodeles, whose development has often been compared with that of temnospondyls (e.g., **Schoch and Carroll, 2003**; **Schoch, 2006**; **Fröbisch et al., 2007**; **Germain and Laurin, 2009**; **Fröbisch et al., 2015**). For instance, **Vorobyeva and Hinchliffe (1996)** documented the larval functional constraints linked to early forelimb use that may cause an early development of manual digits 1 and 2, compared with other tetrapods, as briefly discussed below. However, in the case of our seven cranial characters, there is no evidence of functional constraints. This is a little-investigated topic, but all these bones apparently form a single developmental module of the urodele skull (**Laurin, 2004**). For the appendicular data, functional constraints might explain the more subdued phylogenetic signal, but this will have to be determined by additional research.

The finding that the postcranial characters that we analyzed contain relatively little phylogenetic signal may raise doubts about the claims that have been made about the phylogenetic implications of other such data. Specifically, **Carroll et al. (1999)** stated that the neural arches ossify before the centra in frogs and temnospondyls, but not in salamanders, caecilians or lepospondyls. When it was found that the centra do ossify first in a few cryptobranchoid salamanders, **Carroll (2007, p. 30)** took this as “strong evidence that the most primitive crown-group salamanders had a sequence of vertebral development that is common to frogs and labyrinthodonts [including temnospondyls] (but distinct from that of lepospondyls)”. In fact, apart from tail regeneration in *Hyloplesion* and *Microbrachis* (where the centra ossify before the neural arches: **Olori, 2015**; **Fröbisch et al., 2015**; **Vos et al., 2018**), only one incompletely ossified vertebral column (referred to *Utaherpeton*) is known of any putative lepospondyl. “In this specimen, […] five neural arches […] have ossified behind the most posterior centrum” (**Carroll and Chorn, 1995, 40–41**). Carroll’s **(2007**, **p. 85)** claim that “the centra always ossified prior to the arches” in lepospondyls is therefore rather puzzling.

**Fröbisch et al. (2007)** and **(2015)** pointed out that the first two digital rays (digits, metapodials and distal carpals/tarsals) ossify before the others (“preaxial polarity”) in salamanders and the temnospondyls *Apateon, Micromelerpeton* and *Sclerocephalus*, while the fourth ossifies first (“postaxial polarity”) in amniotes, frogs and “probably” (**Fröbisch et al., 2015, pp. 233, 234**) the lepospondyls *Microbrachis* and *Hyloplesion*. This latter inference, however, is based only on a delay in the ossification of the fifth ray that is shared specifically with sauropsid amniotes (**Olori, 2015**). Ossification sequences (however partial) of the other four rays in any lepospondyl are currently limited to the tarsus of *Batropetes*, which clearly shows preaxial polarity (**Glienke, 2015, fig. 6O–S**; **Marjanović and Laurin, 2019**), and that of the putative (but see **Clack et al., 2019**) lepospondyl *Sauropleura*, in which likewise the second distal tarsal ossified before all others (**Marjanović and Laurin, 2019**). Outside of temno- and lepospondyls, **Marjanović and Laurin (2013, 2019)** presented evidence that preaxial polarity is plesiomorphic, widespread and dependent on the use of the still developing limbs for locomotion, which would explain why it was independently lost in amniotes and frogs and reduced (the second ray still forms first, but the delays between the rays are much reduced so that all form nearly at the same time) in direct-developing salamanders as well as in the limb regeneration of terrestrial postmetamorphic salamanders (**Kumar et al., 2015**). It may be relevant here that the PH2 (**Fig. 1f**), favored by our appendicular data, groups exactly those sampled taxa in a clade that are known to have preaxial polarity in limb development. To sum up, neither our own analyses nor the previous works that we cited above demonstrated conclusively that ossification sequences of postcranial elements provide reliable clues about the origin of extant amphibians.

In contrast, we are reasonably confident about our results on the cranial ossification sequences. Given the phylogenetic signal we have found in our cranial datasets, we think that ossification sequence data should eventually be added to phenotypic datasets for analyses of tetrapod phylogeny. Indeed, an analysis of amniote phylogeny using data from organogenesis sequences (coded using event-pairing in Parsimov) already exists (**Werneburg and Sánchez-Villagra, 2009**). The usefulness of such data for phylogenetic inference was further tested, with encouraging results, by **Laurin and Germain (2011)**, and the present analysis adds additional support for it.

### Indirect support for the lepospondyl hypothesis from temnospondyls

The strong support for the lepospondyl hypothesis that we have found in cranial data is surprising because cranial ossification sequence data, especially those of the Permo-Carboniferous temnospondyl *Apateon*, have often been claimed to contradict the LH (lepospondyl hypothesis, **Fig. 1d**). Similarities between *Apateon* and extant urodeles, in particular the supposedly “primitive” hynobiid *Ranodon*, have often been emphasized (**Schoch and Carroll, 2003**; **Schoch and Milner, 2004**; **Carroll, 2007**; **Schoch, 2014a**). However, other studies have already raised doubts about some of these claims (e.g., **Schoch, 2006**; **Anderson, 2007**; **Germain and Laurin, 2009**). **Schoch (2006)** and **Anderson (2007)** concluded that most characters shared between *Apateon* and urodeles were plesiomorphies. **Germain and Laurin (2009)** also demonstrated that, far from being very similar to the ancestral urodele morphotype (contra **Schoch and Carroll, 2003** or **Carroll, 2007**), the cranial ossification sequence of *Apateon* was statistically significantly different from that of the hypothetical last common ancestor of all urodeles (as suspected by **Anderson, 2007**). However, these earlier studies did not clearly show which of the various hypotheses on lissamphibian origins the ossification sequences of *Apateon* spp. – or the newly available partial sequence (**Werneburg, 2018**) of the phylogenetically distant temnospondyl *Sclerocephalus* – supported most. This is what we have attempted to do here.

Unfortunately, the development of lepospondyls is too poorly documented to be incorporated into the cranial analyses, but we included two lepospondyls in analyses of appendicular data. These analyses weakly favor a polyphyletic origin of extant amphibians, with both temno- and lepospondyls in the amphibian clade, a hypothesis that has not been advocated seriously for decades (**Milner, 1993, fig. 5B**) as far as we know. However, given the moderate phylogenetic signal in these data, we view these results with skepticism. **Olori (2011)**, using event-pairing with Parsimov (**Jeffery et al., 2005**) and PGi (**Harrison and Larsson, 2008**), analyzed lepospondyl postcranial ossification sequences and concluded that support for the three hypotheses that she tested (TH/DH with two different positions for *Micromelerpeton*, and LH) did not differ significantly. By contrast, our analyses of the postcranial data indicate a stronger support for polyphyly (PH2) than for the TH/DH, which is only a distant second (**Table 8**) or third (behind PH2 and LH; **Table 9**) depending on the analyses. **Olori (2011)** performed no statistical test of phylogenetic signal of her data, though a related test (performing phylogenetic analyses on the data) yielded trees (**Olori, 2011, fig. 5.5–5.7**) that are largely incongruent with the established consensus, in which most large taxa (Mammalia, Testudines, Lissamphibia, etc.) are para- or polyphyletic. Olori’s **(2011)** results, like ours, support the conclusion that the phylogenetic signal in postcranial ossification sequence data is low.

Given the current limitations in the availability of developmental data in Paleozoic stegocephalians, we hope to have demonstrated that cranial ossification sequences of amniotes, lissamphibians and temnospondyls provide support for the LH that is independent of the phylogenetic analyses of **Laurin (1998), Pawley (2006, appendix 6)** or **Marjanović and Laurin (2009, 2019)**. This independence is important because the cranial ossification sequence data cannot rival the morphological data in terms of data availability, simply because growth sequences of extinct taxa are rare (**Sánchez-Villagra, 2012**), but having a fairly independent line of evidence to investigate a major evolutionary problem is clearly advantageous. We hope that the modest methodological progress made in this study will stimulate the search for fossilized ontogenies (**Cloutier, 2010**; **Sánchez-Villagra, 2010**).

## Supporting information

Supplements. Data matrices and trees

## Acknowledgements

Jennifer Olori, two anonymous reviewers and the Recommender Robert Asher made helpful comments that improved this paper. D. M. would further like to thank Ralf Werneburg for an electronic reprint of his 2018 paper, Nadia Fröbisch for discussion of limb development in salamanders, and Daniel Field for discussion of molecular divergence times and the fossil record.

## Additional information

### Author contributions

ML designed the study, supervised the data collection, analyzed the data and wrote much of the draft; OL collected most of the ossification sequence data; DM added data to the database (mostly of Paleozoic taxa), updated the timetrees, participated in the writing, and drafted Figure 1.

### Data availability

All data used in this study are available as Supplementary material (see below)

### Supplementary information

Data matrices and trees used in this analysis can be downloaded in NEXUS format for Mesquite from: https://www.biorxiv.org/content/10.1101/352609v3.supplementary-material.

## Appendix

### Sources of data for ossification sequences

Empty cells indicate that these data are unavailable. Three methods were examined, and we used the one for which most data were available (position in the ossification sequence, last column).

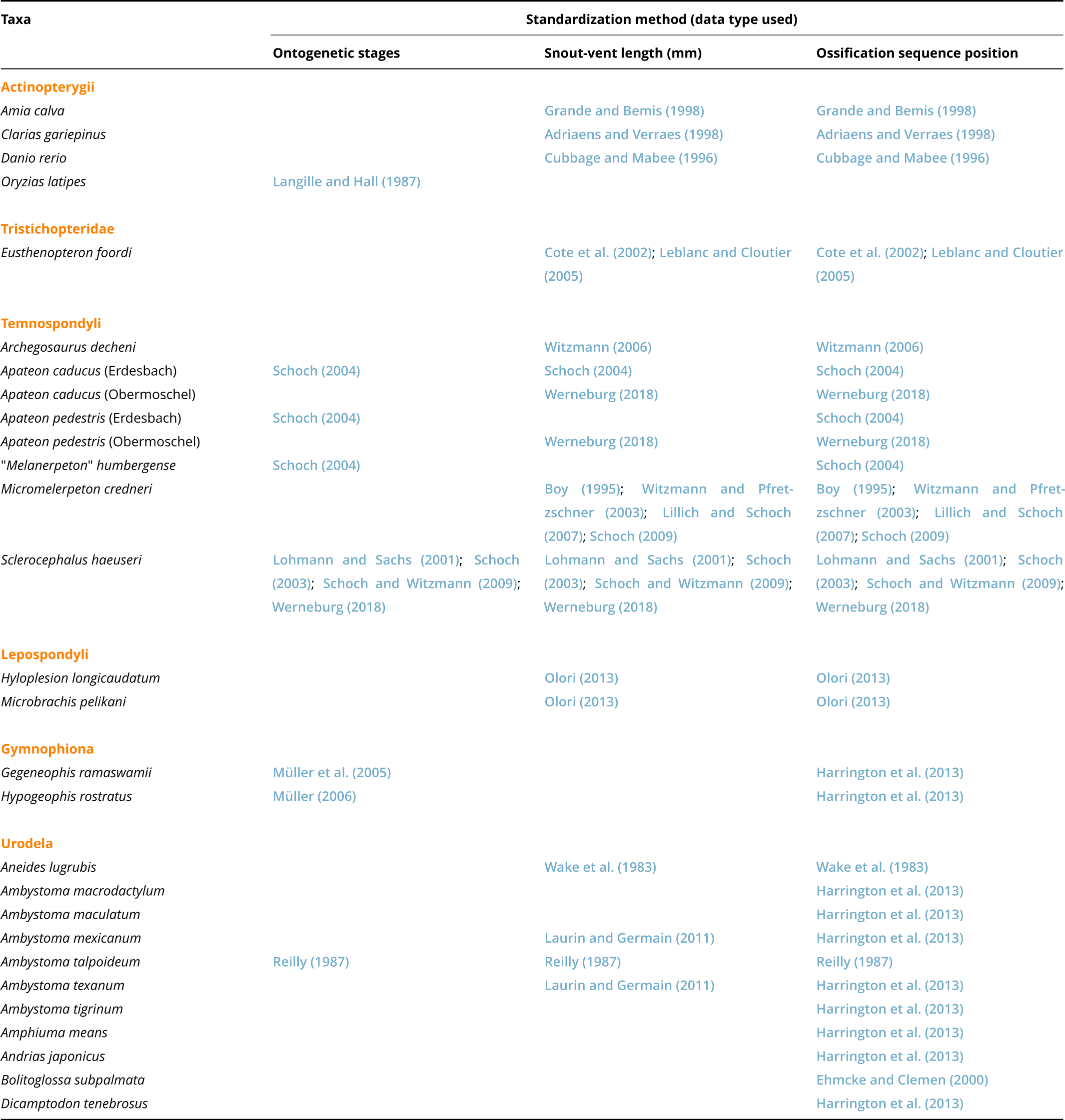
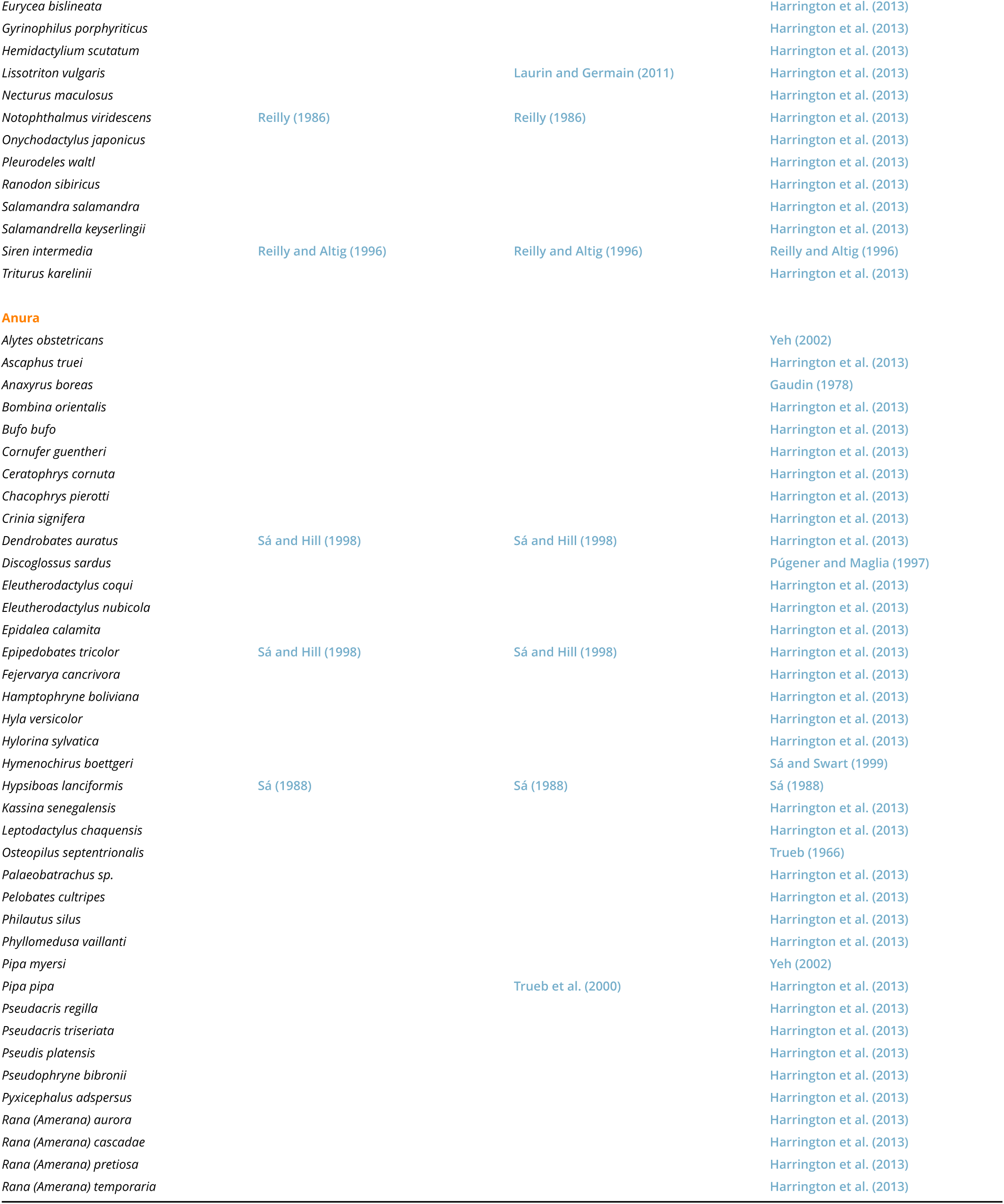
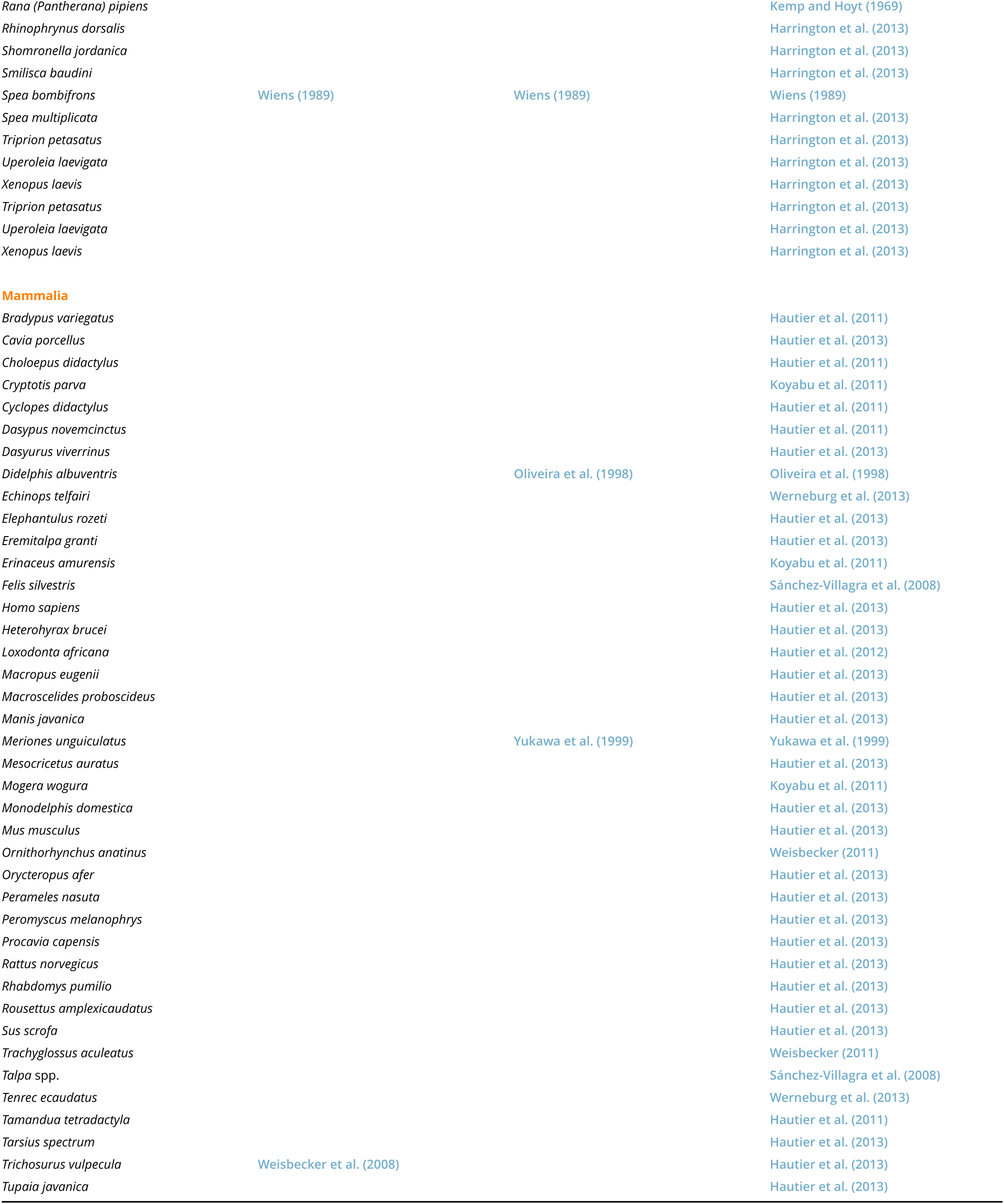
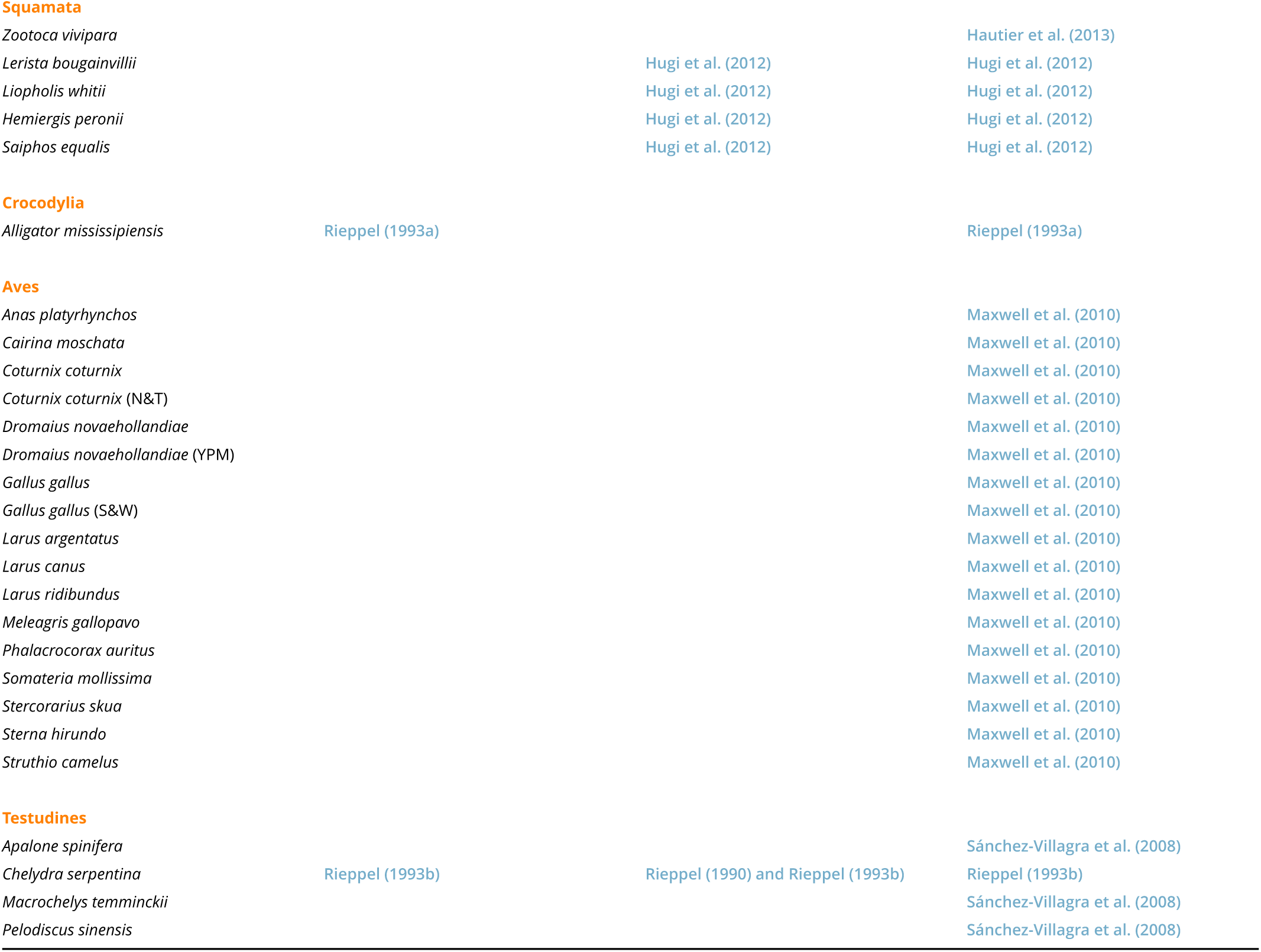

